# Early alterations of social brain networks in young children with autism

**DOI:** 10.1101/180703

**Authors:** Holger Franz Sperdin, Ana Coito, Nada Kojovic, Tonia Rihs, Reem Kais Jan, Martina Franchini, Gijs Plomp, Serge Vulliémoz, Stéphan Eliez, Christoph Martin Michel, Marie Schaer

## Abstract

Social impairments are a hallmark of Autism Spectrum Disorders (ASD), but empirical evidence for early brain network alterations in response to social stimuli is scant in ASD. Here, we recorded the gaze patterns and brain activity of toddlers and preschoolers with ASD and their typically developing (TD) peers while they explored dynamic social scenes. Source-space directed functional connectivity analyses revealed the presence of network alterations in the theta frequency band, manifesting as increased driving (hyper-activity) and stronger connections (hyper-connectivity) from key nodes of the social brain associated with autism. Further analyses of brain-behavioural relationships within the ASD group suggested that compensatory mechanisms from dorsomedial frontal, inferior temporal and insular cortical regions were associated with lower clinical impairment and less atypical gaze patterns. Our results provide strong evidence that directed functional connectivity alterations of social brain networks is a core component of atypical brain development at early stages of ASD.

---

Early preferential attention to social cues is a fundamental mechanism that facilitates interactions with other human beings. During the third trimester of gestation, the human fetus is already sensitive to both voices (Decasper & Spence, 1986) and face-like stimuli (Reid et al., 2017). Newborns orient to biological motion (Simion et al., 2008) and prefer their mothers’ voices to those of other females (Decasper & Fifer, 1980). Infants as young as two weeks imitate faces and human gestures (Meltzoff & Moore, 1977). The orientation to and interaction with social cues during infancy drives the later acquisition of social communication skills during toddler and preschool years. As function of experience, the repeated exposure leads to the progressive emergence of adaptive interactions with conspecifics. Alongside, the brain develops a network of cerebral regions specialized in understanding the social behaviors of others. This network includes the orbitofrontal and medial prefrontal cortices, the superior temporal cortex, the temporal poles, the amygdala, the precuneus, the temporo-parietal junction, the anterior cingulate cortex (ACC) and the insula (Brothers, 1990; Frith & Frith, 2010; Adolphs, 2009; Blake-more, 2008). Collectively, these areas form the *social brain* and are all implicated to some extent in processing social cues and encoding human social behaviors (Brothers, 1990; Frith & Frith, 2010; Adolphs, 2009; Blakemore, 2008).

Autism is a life-long lasting, highly prevalent neurodevelopmental disorder that affects core areas of cognitive and adaptive function, communication and social interactions (Christensen et al., 2016). A common observation in infants later diagnosed with ASD is the presence of less sensitivity and diminished preferential attention to social cues during the first year of life (Osterling & Dawson, 1994). Toddlers with ASD orient preferentially to non-social contingencies (Klin et al., 2009). Indifference to voices (Sperdin & Schaer, 2016) and faces (Grelotti et al., 2002) in ASD leads to deficits in the development of adapted social interactions with others and to difficulties in understanding human behaviors. It is not established why children with ASD show insensitivity to stimuli with social contingencies at early developmental stages, but this apparent indifference to social cues ultimately hinders the normal development of the *social brain* network or parts thereof (Pelphrey et al., 2011; Gotts et al., 2012). Some authors propose that deficits in the development of social cognition (such as learning to attribute mental states to others, “theory of mind “ (Frith, 1989)) and/or in sensory processing (Dinstein et al., 2012) prevent children with ASD to actively and appropriately engage with social stimuli. Another hypothesis suggests that they have difficulties building up stimulus-reward contingencies for social stimuli, due to a reduced motivation to attend and engage with them. Regardless of the reasons behind reduced social orienting, diminished interaction and exposure to social stimuli may in turn impede the development of the *social brain* at early developmental stages in ASD (Chevallier et al., 2012; Dawson et al., 2004).

Evidence remains limited for brain network alterations in response to socially meaningful stimuli in ASD during the period spanning the toddler to preschool years, partly because the acquisition of data during that age period is extremely challenging (Raschle et al., 2012). However, studying very young children with ASD, closer to their diagnosis, is all the more important when recent findings suggest the presence of major developmental changes in large-scale brain networks between adults and younger individuals with ASD (Nomi & Uddin, 2015). Currently, it remains unclear how autism affects the development of the functional brain networks implicated in the processing of socially meaningful information at early developmental stages. A better delineation of the timing and nature of the neurodevelopmental alterations associated with core social deficits in autism may in turn help to improve therapeutic strategies.

Electroencephalography (EEG) is as a powerful noninvasive method to study atypical brain responses to social stimuli in clinical pediatric populations with ASD. For example, surface-based experiments have reported aberrant evoked potentials in response to dynamic eye gaze in infants at high-risk for ASD (Elsabbagh et al., 2012) or to speech stimuli in toddlers with ASD (Kuhl et al., 2013) with differences in resting EEG power in infants at high-risk for ASD (Tierney et al., 2012). Whilst useful, most of the EEG experiments performed on very young children with ASD (younger than four years) have been done with few electrodes only and the analysis restricted to the sensor space. Therefore, hypothetical alterations in the functional brain networks underlying the processing of social stimuli remain to be determined for that age period in ASD.

Here, we recorded high-density EEG and high resolution eye-tracking in toddlers and preschoolers with ASD and their TD peers as they watched naturalistic and ecologically valid dynamic social movies. Using data-driven methods, we first investigated whether the visual exploration behavior was atypical in toddlers and preschoolers with ASD using kernel density distribution estimations. Then, we explored whether their ongoing source-space directed functional connectivity was altered compared to their TD peers using Granger-causal modeling applied to the EEG source signals. This method estimates brain connectivity in the frequency domain. It identifies which brain regions are the key drivers of information flow in a brain network and directional relationships between brain regions that belong to a network (Coito et al., 2016b). Finally, we looked for relationships between directed functional connectivity measures, visual exploration behavior and clinical phenotype. As toddlers with ASD have less preferential attention for social cues, we hypothesized that they would show both a different visual exploration behavior of the dynamic social images and altered directed functional connectivity patterns in brain regions involved in processing social information compared to their TD peers.

## Results

### Summed outflow

The summed outflow (i.e.- the amount of information transfer) is a measure that reflects the importance (i.e.-the amount of driving) of a given region of interest (ROI) in the network (see method section). To understand the functional wiring and the dynamic flow underlying the processing of the dynamic social stimuli, we used a data-driven method to explore in which frequency band the highest summed outflow occurred in 82 ROIs across the whole brain. A ROI with a strong summed outflow has a key role in directing the activity towards other ROIs in the network. The strongest summed outflow across the whole brain occurred in the theta band (4 - 7*Hz*) in both groups. The summed outflow of the largest drivers across frequencies is illustrated for each group in Figure 1a. As can be seen the largest peaks of activity are present in the theta band range (4 - 7*Hz*) in both group. The global driving in the theta band did not differ between the groups (*t* = 0.6201*, p* = 0.536). In addition, several regions common to both groups showed a large driving (summed outflow) in this theta band, and notably the bilateral medial frontal and superior orbitofrontal regions, the bilateral hippocampi, the bilateral ACC and the right amygdala (Figure 1b).

**Figure 1.**
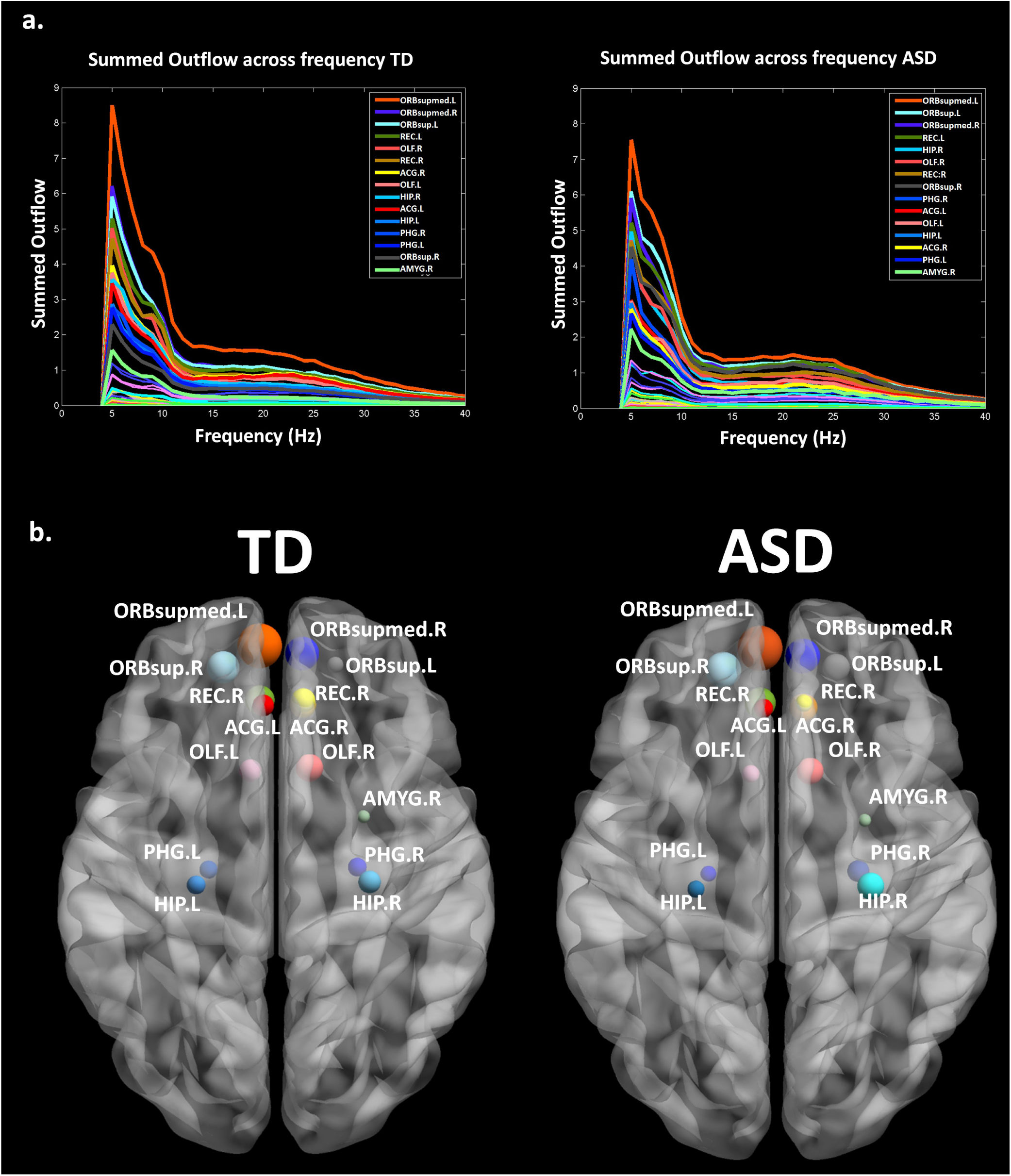
a. The summed outflow of the largest drivers across frequencies is illustrated for each group (TD, Left; ASD, Right).b. Regions consistently showing large driving in both groups (TD, Left; ASD, Right): the bilateral superior frontal gyri, medial orbital (ORBsupmed.L, ORBsupmed.R), the bilateral superior frontal gyri, orbital (ORBsup.L, ORBsup.R), the bilateral rectus gyri (REC.L, REC.R), the bilateral olfactory cortices (OLF.L,OLF.R), the bilateral hippocampi (HIP.L, HIP.R), the bilateral parahippocampi (PHG.L, PHG,R), the bilateral anterior parts of the cingulate gyri (ACG.L, ACG.R) and the right amygdala (AMYG.R). Summed outflows are represented as spheres: the larger the sphere, the higher the summed outflow.

Thereafter, we characterized the differences in the summed outflow in the theta band across all brain regions between the groups. Amongst all the ROIs, we identified six ROIs that showed a statistically higher driving (stronger summed outflow) in the ASD group in comparison to the TD group (*Mann - W hitney - Wilcoxontest,two - tailed, p <* 0.05): the right orbital part of the superior frontal gyrus (*W s* = 267*,z* = −2.088*.p* = 0.037*,r* = −0.348), the bilateral orbital parts of the middle frontal gyri (*Le f t* : *W s* = 259*,z* = −2.341*, p* = 0.019*,r* = −0.39;*Right* : *W s* = 252*,z* = −2.563*, p* = 0.01*,r* = −0.427), the right middle cingulate gyrus (*W s* = 259*,z* = −2.341*, p* = 0.019*,r* = −0.390), the left superior occipital gyrus (*W s* = 270*,z* = −1.993*, p* = 0.047*,r* = −0.332), and the left superior temporal gyrus (STG) (*W s* = 255*,z* = −2.468*, p* = 0.013*,r* = −0.411) (Figure 2). This indicates the presence of a stronger driving from these regions in the toddlers and preschoolers with ASD compared to their TD peers when viewing the dynamic social images. No other regions had a higher driving in the TD group compared to the ASD group.

**Figure 2.**
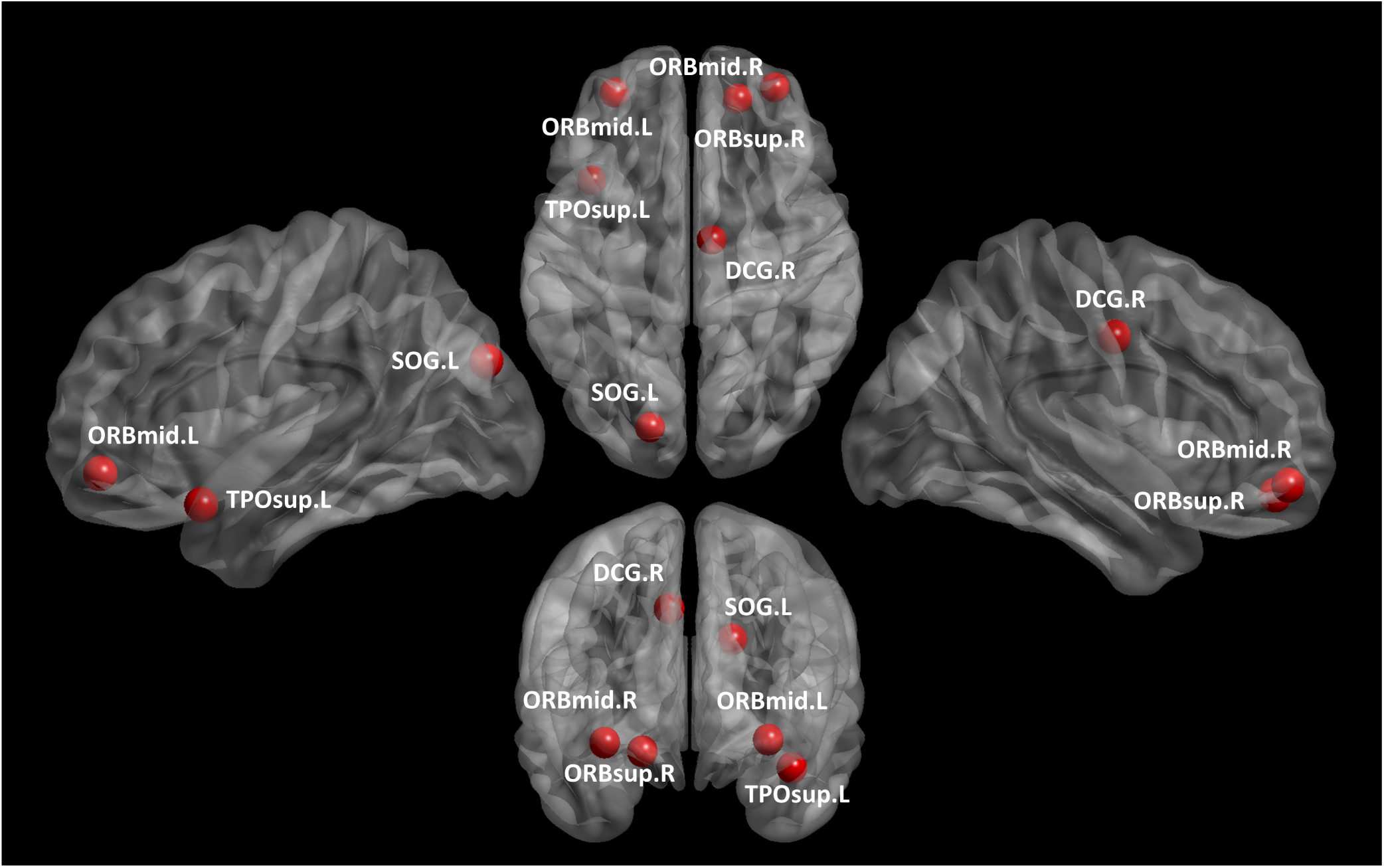
The 6 ROIs with a statistically significant increased summed outflow in the ASD group compared to their TD peers: the right orbital part of the superior frontal gyrus (ORBsup.R), the bilateral orbital parts of the middle frontal gyri (ORBmid.L, ORBmid.R), the right middle cingulate gyrus (DCG.R), the left superior occipital gyrus (SOG.L), and the left superior temporal gyrus (TPOsup.L).

### Region-to-region directed functional connectivity

We looked for differences in the region-to region directed functional connectivity using Granger-causal modelling (see method section) from each of the six nodes separately in both groups. The results indicated that all the connections in the toddlers and preschoolers with ASD were stronger than the strongest connections in the TD participants (*Mann* – *W hitney* – *Wilcoxon, two* – *tailed*, *p* < *0.05*, *Ben jamini - Hochberg* = 0.05). This suggests the presence of hyper-connectivity in the toddlers and preschoolers with ASD. In both groups, the strongest connections was from a node within the frontal cortex: the right superior orbito-frontal region (OFC). The region-to-region directed functional connectivity from the right superior OFC is illustrated in Figure 3. The estimation of the region-to-region directed connectivity (i.e. to which other ROIs the activity was directed) revealed major differences between the groups. In the TD group, the right superior OFC had directed connections towards the left middle frontal, the right precentral, the left superior frontal, the left supramarginal, the left cuneus, left hippocampus, right paracentral, right opercular part of the inferior frontal cortex, left inferior parietal cortex and the right angular gyrus. In the group of toddlers and preschoolers with ASD, the right superior OFC drove information flow towards the left orbital part of the medial frontal cortex, left amygdala, left supramarginal, right supplementary motor area, right lingual gyrus, right medial part of the medial frontal cortex, left paracentral, left inferior frontal triangularis region, right hippocampus and right orbital part of the inferior frontal cortex. Thus, the analysis reveals a different network pattern in the toddlers and preschoolers with ASD compared to their TD peers. We provide the directed connections results from the five remaining ROIs in the supplementary information.

**Figure 3.**
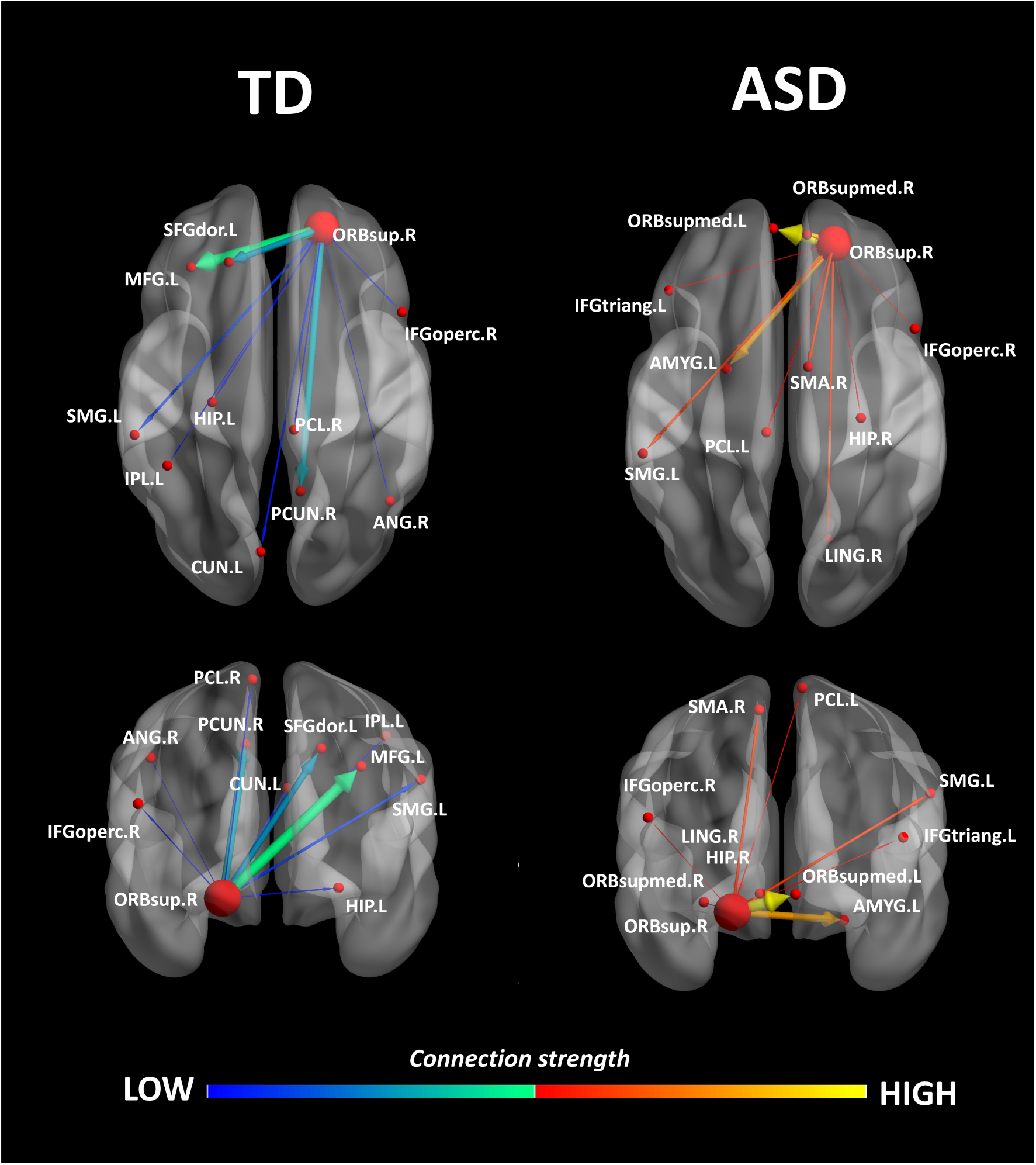
Region-to-region directed functional connectivity in TD group (left) and in ASD group (right). Connections strength and networks are different between the groups. Exemplary node of the right orbital part of the superior frontal gyrus (ORBsup.R) (in large red). For the TD group, the ORBsup.R drove activity towards the left Superior frontal gyrus, dorsolateral (SFGdor.L), the left middle frontal gyrus (MFG.L), the right inferior frontal gyrus, opercular (IFGoperc.R), the left hippocampus (HIP.L), the left cuneus (CUN.L), the left inferior parietal gyrus (IPL.L), the left supramarginal gyrus (SMG.L), the right angular gyrus (ANG.R), the right precuneus (PCUN.R) and the right paracentral lobule (PCL.R). In the ASD group, the node is driving to the left inferior frontal gyrus, triangular (IFGtriang.L), the right inferior frontal gyrus, orbital (ORBinf.R), the right supplementary motor area (SMA.R), the bilateral superior frontal gyrus, medial orbital (ORBsupmed.L ORBsupmed.R), the right hippocampus (HIP.R), the left amygdala (AMYG.L), the right lingual gyrus (LING.R), the supramarginal gyrus (SMG.L) and the paracentral lobule (PCL.L).

### Correlations with ADOS-2, PEP-3, VABS-II scores and gaze Proximity Index

We further explored associations between the summed outflow from the ROIs and clinical and behavioural phenotypes. None of the correlations between the summed outflow and ADOS-2 severity scores survived False discovery rate (FDR) correction (*Ben jamini - Hochberg* = 0.05). However, we found strong positive correlations between the summed outflow in the right lingual area and scores from the socialization domain (*rs* = 0.751*, p* = 0.0003*,two - tailed, <* 0.05;*Ben jamini - Hochberg* = 0.05) as well as with scores from the leisure and play skills subdomain of the VABS-II (*rs* = 0.802*, p* = 0.0001*,two - tailed, <* 0.05;*Ben jamini - Hochberg* = 0.05). Higher summed outflow within the left Heschl area near the posterior convolutions of the insula and the left rolandic operculum near the circular sulcus of the insula rostrally were positively related to better fine (*rs* = 0.745*, p* = 0.0004*,two - tailed, <* 0.05;*Ben jamini - Hochberg* = 0.05) and gross motor skills (*rs* = 0.744, *p* = 0.0004, *two - tailed, <* 0.05;*Ben jamini - Hochberg* = 0.05) as measured by the PEP-3. Finally, toddlers and preschoolers with ASD with a gaze pattern similar to their TD peers showed an increased driving within the left middle cingulate cortex (*rs* = 0.726*, p* = 0.0007*,two-tailed, <* 0.05;*Ben jamini - Hochberg* = 0.05) and the right paracentral lobule (*rs* = 0.738*, p* = 0.0005*,two - tailed, <* 0.05;*Ben jamini - Hochberg* = 0.05). The correlations between the summed outflows and the Proximity Index (See method section), VABS-II scores and PEP-3 scores are displayed in Figure 4.

**Figure 4.**
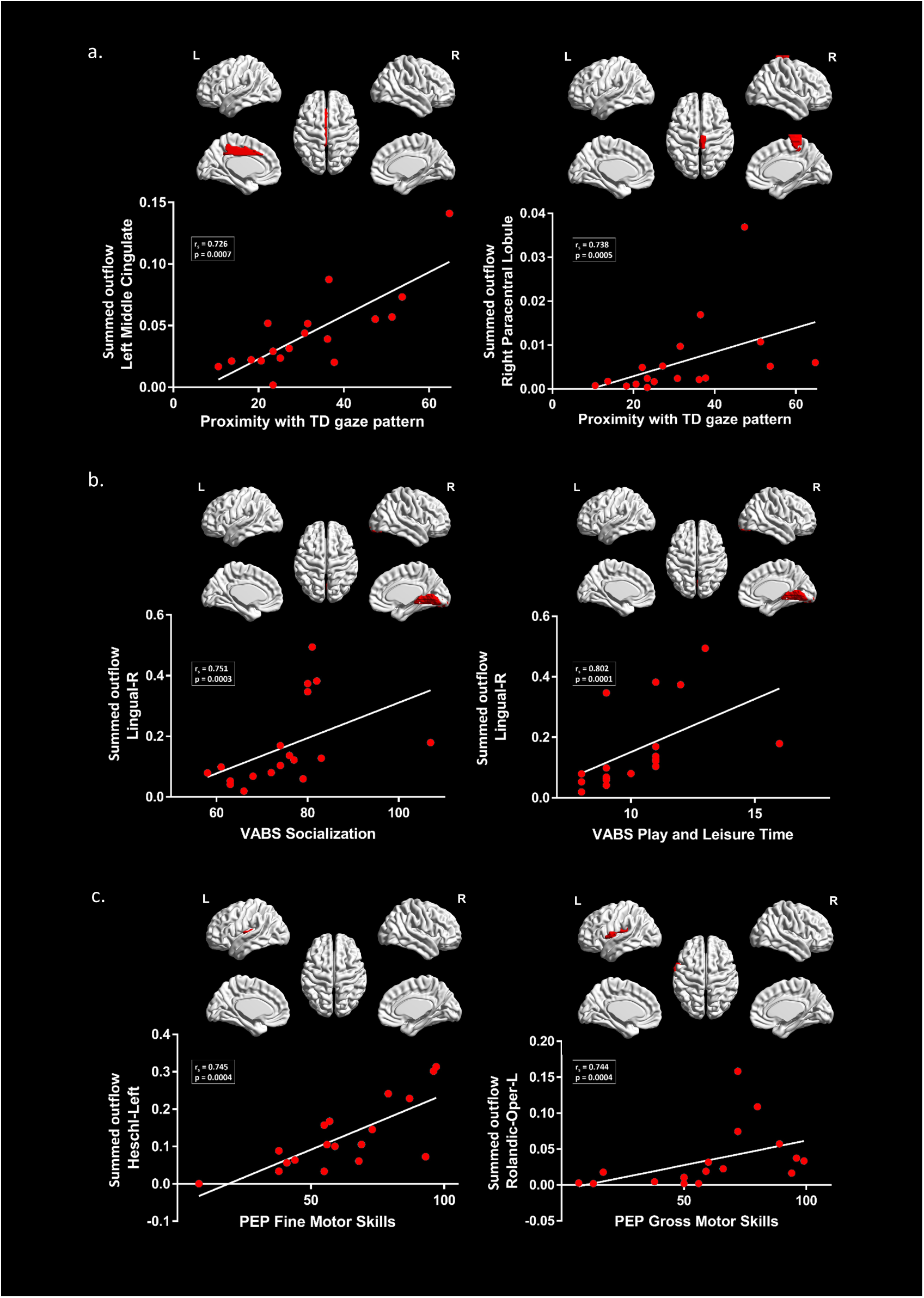
Correlations between the summed outflows and (a.) Proximity Index (b.) VABS-II scores and (c.) PEP-3 scores.

## Discussion

Abnormal processing of social cues is a hallmark of ASD (Chevallier et al., 2012; Dawson et al., 2004; Dichter et al., 2009; Elsabbagh et al., 2012; Gotts et al., 2012; Greene et al., 2011; Klin et al., 2009; Pelphrey et al., 2011). However, evidence for alterations of social brain networks at early stages of ASD is scant. Using data-driven methods, we observed aberrant gaze patterns together with altered directed functional connectivity in toddlers and preschoolers with ASD when exploring dynamic social stimuli compared to their TD peers. These differences manifested as increased driving and hyper-connectivity in the theta frequency band from nodes that includes the right orbital part of the superior frontal gyrus, the bilateral orbital parts of the middle frontal gyri, the right middle cingulate gyrus, the left superior occipital gyrus and the left STG. To the best of our knowledge, this is the first evidence indicating concomitant alterations in the visual exploration of dynamic social images and in the directed functional connectivity involving key nodes of the *social brain* (Brothers, 1990; Frith & Frith, 2010; Adolphs, 2009; Blakemore, 2008) at early stages of ASD.

Our results indicate that the highest information transfer (i.e. summed outflow) occurred at the global brain level in the theta frequency band (4 - 7*Hz*). Slow waves are prominent brain rhythms during infancy and toddlerhood. Throughout development, they modulate attentional brain states, encode specific information and ease communication between neuronal populations. Theta band activity is associated with sustained anticipatory attention, memory encoding, emotional processing and cognitive performance during infancy, toddlerhood and preschool years (Saby & Marshall, 2012). EEG experiments in individuals with ASD show a reduction or an increase in the coherence patterns in the theta frequency band compared to their TD peers at various ages and under different experimental conditions with sensor-space based analysis (Schwartz et al., 2016). In very young children, theta band activity is linked to the development of the *social brain.* For example, surface-based EEG experiments in TD infants report a greater theta power to social versus non-social stimuli at 12 months (Jones et al., 2015). Theta increases during attention to social stimulation in infants and preschool aged children (Orekhova et al., 2006). Hence, social contingencies modulate theta band activity. Similarly, our results show high information transfer in the theta band when toddlers and preschoolers explore dynamic social stimuli. However, most of the available EEG experiments and analysis thereof are restricted to the scalp surface. Therefore, information is limited regarding the involvement of specific brain regions when young children are exposed to social stimuli. Here, our data-driven source-space approach that estimates brain connectivity in the frequency domain revealed not only high information transfer in the theta frequency band, but also, the involvement of the bilateral medial and the superior orbital frontal regions, the bilateral hippocampi, the bilateral ACC and right amygdala. These areas are implicated in processing social cues and encoding human social behaviours (Brothers, 1990; Frith & Frith, 2010; Adolphs, 2009; Blakemore, 2008).

Our results further indicate the presence of an altered theta response (higher driving) in the toddlers and preschoolers with ASD compared to their TD peers within key regions of the *social brain*. In TD individuals, theta generates within the frontal cortices or the ACC (Asada et al., 1999). In comparison to their TD peers, both areas develop differently in young toddlers later diagnosed with ASD (Schumann et al., 2010). Accordingly, our results raise the possibility that the brain regions generating theta also follow a different development as driving in the theta band from frontal and cingulate regions was altered in our participants with ASD compared to their TD peers. A magnetoencephalography (MEG) study using a source-space approach reported lower coherence in the theta band within parietal and occipital regions but their sample included adolescents with ASD only (Ye et al., 2014). The differences between this specific study and our could stem from either variations in the age groups (adolescents versus toddlers and preschoolers), the stimuli used (grey cross inside a white circle versus dynamic biological visual stimuli) or the methods. More generally, several factors explain discrepancies in brain connectivity results between studies. The type of connectivity measures applied, the approach (surface versus source based), the brain regions analysed and frequency bands examined are variables that influence the results (Mohammad-Rezazadeh et al., 2016). In ASD, hyper-connectivity is prevalent in younger populations (that is, infants at high-risk for ASD, toddlers and preschoolers with ASD) while hypo-connectivity is more observed during adolescence and adulthood in ASD (Uddin et al., 2013b). Conversely, a developmental shift occurs in brain growth with an initial period of early brain overgrowth followed by normalization sometime during adolescence (Courchesne et al., 2011). Accordingly, structural white matter connectivity studies also highlight this shift from higher structural connectivity in very young children with ASD to lower connectivity in older children with ASD (Hoppenbrouwers et al., 2014; Conti et al., 2015). Therefore, higher-driving and hyper-connectivity from key nodes of *the social brain* in the theta frequency band in our ASD sample is consistent with reports in the literature when considering the very young age of our participants (around 3 years of age on average).

We found increased activity from nodes of the orbital and medial parts of the frontal cortex and cingulate cortex in the toddlers and preschoolers with ASD. Both areas have been implicated in various complex aspects of social cognition, social reward, social perception and social behaviour (Jonker et al., 2015; Apps et al., 2013). Metabolic changes within the medial prefrontal cortex and the cingulate cortex are correlated with social interaction impairments in childhood ASD (Ohnishi et al., 2000). Several experiments report structural (Patriquin et al., 2016) and functional (Gotts et al., 2012; Patriquin et al., 2016; Greene et al., 2011; Cheng et al., 2015) alterations within these brain areas in school aged children, adolescents and adults with ASD when compared to their TD peers. A recent study described hyper-connectivity within the ACC and bilateral insular cortices in a sample including children aged between seven to 12 years (Uddin et al., 2013a). Some authors propose that the two together form the *salience network*, whose role is to direct attention to behaviourally-relevant stimuli (Menon & Uddin, 2010). Although we didn’t find differences in the driving from the insula compared to the TD peers, there is an increasing number of evidence showing an abnormal development of the *salience network* or components thereof in ASD (Uddin, 2015), which may partially explain the limited interest for and engagement with social stimuli that is often observed in individuals with ASD and that constitutes a hallmark of the disorder (Klin et al., 2009; Pelphrey et al., 2011). Accordingly, the toddlers and preschoolers with ASD had a different visual exploration behaviour of the dynamic social stimuli raising the possibility of a reduced interest to visually engage with them. Alternatively, alterations in driving from these regions could partially reflect a reduced motivation to attend and engage with the dynamic social stimuli (Chevallier et al., 2012; Dawson et al., 2004).

We also found higher driving from the superior temporal and occipital gyri. Those brain areas are implicated in the processing of biological motion, in analysing the intentions of other people’s actions and self-reflection (Pelphrey & Carter, 2008; Pelphrey et al., 2005; Pelphrey & Morris, 2006; Pelphrey et al., 2004). Our result would suggest that the exploration of the dynamic social visual stimuli that contained biological movements led to altered driving from these brain areas in toddlers and preschoolers with ASD compared to their TD peers. Beyond functional and structural brain alterations reported elsewhere in older children and adults with ASD, our results suggest for the first time, the presence of alterations in the driving of information flow from brain areas implicated in social information processing during the viewing of naturalistic dynamic social images in toddlers and preschool with ASD.

We further explored associations between the driving in the nodes of the network (that is, the summed outflow) and clinical and behavioural phenotypes. We didn’t observe any significant relationships between summed outflow and the ADOS severity scores after FDR correction. Notwithstanding, we observed an increased driving from the median cingulate cortex and the paracentral lobule in the toddlers and preschoolers with ASD who had a more similar visual exploration pattern to their TD peers. Thus, an improved visual exploration pattern of the dynamic social images was related to increased summed outflow from these regions. Likewise, higher summed outflow from the right lingual area was related to better socialization behaviour and leisure and play skills. Higher summed outflow from the left Heschl’s area near the posterior convolutions of the insula and the left rolandic operculum near the circular sulcus of the insula rostrally were positively related to better fine and gross motor skills. As such, increased activity within dorsomedial frontal, inferior temporal and insular cortical regions were associated with lower clinical impairment and less atypical gaze patterns. The presence of hyperactivity within relevant brain region has been interpreted as a possible compensatory mechanism when performing a social target detection task, in adults with ASD at least (Dichter et al., 2009). We speculate that the hyper-driving from these brain regions might be a mechanism to compensate for atypical development of the brain’s circuitry over time as higher directed functional connectivity was related to better visual exploration of dynamic social images, improved socialization and motor behaviours.

The collective results suggest that directed functional connectivity network alterations within regions of the *social brain* are present at early stages of ASD, justifying further investigation into how early therapeutic interventions targeting social orienting skills may help to remediate social brain development during this critical age period when plasticity is still possible.

## Methods

### Participants

Recruitment of toddlers and preschoolers with ASD was achieved via clinical centres specialized in ASD and French-speaking parent associations. TD toddlers and preschoolers were recruited via announcements in the Geneva community. Prior to the experiments, all the procedures were approved by the Ethics Committee of the Faculty of Medicine of the University of Geneva Hospital in accordance with the ethical standards proclaimed in the Declaration of Helsinki. For all participants, an interview over the phone and a medical developmental history questionnaire were completed before their initial visit. All participants’ parents gave their informed consent prior to inclusion in the study. 120 participants were recruited for the experiment. We did not manage to put the EEG cap on the head of 23 ASD and 7 TD participants. We managed to put the cap on 90 participants. Out of those, we excluded 28 ASD and 26 TD participants because of too many movement-related artefacts, unrepairable noisy signal, lack of interest, or insufficient amounts of epochs available for subsequent analysis. This was to be expected given the extremely sensitive population at study here. As a result, 36 participants were included in the final sample: 18 young children with ASD (2 females; mean age 3.1 years +/- 0.8, range 2.2-4.4) and 18 age matched (*t* = 2.72*, p* = 0.852) TD peers (5 females; mean age 3.1 years +/- 0.9, range 2.0-4.8). All participants with ASD included in the study received a clinical diagnosis prior to their inclusion in the research protocol. Diagnosis of ASD was rigorously verified and confirmed with either the Autism Diagnosis Observation Schedule – Generic (Lord et al., 2000) or the Autism Diagnosis Observation Schedule, second edition (ADOS-2)(Luyster et al., 2009). The latter contains a toddler module that defines concern for ASD. ADOS assessments were administered and scored by experienced clinicians working at the institution and specialized in ASD identification. In order to compare scores from different modules, we transformed the ADOS-G scores into Calibrated Severity Scores (ADOS-CSS) (Gotham et al., 2009). For the participants that underwent the ADOS-2-toddler module, we calibrated the scores into Severity Scores (Esler et al., 2015). Five children under 30 months of age performed the toddler module of the ADOS-2. All scored in the moderate to severe range of concern for ASD. For all the participants younger than 3 years of age (n=10) at the EEG acquisition, clinical diagnosis was confirmed after one year by a clinician specialized in ASD identification using the ADOS-G or ADOS-2. The mean global ADOS-CSS for the entire group of patients with ASD was 7.9 (SD = 1.6). The assessment of the participants with ASD also included the administration of additional clinical standardized tests. Adaptive behaviour was assessed using the Vineland Adaptive Behaviour Scale-II (VABS-II)(Sparrow et al., 2005), a standardized parent report interview. Developmental level was assessed with the Psycho-educational Profile Third Edition (PEP-3)(Schopler et al., 2005). See Table 1 for characteristics of study participants. Prior to their inclusion in our research protocol, potential TD participants were initially screened for neurological/psychiatric problems and learning disabilities using a medical and developmental history questionnaire before their visit. Moreover, they underwent ADOS-G or ADOS-2 evaluations to exclude any ASD symptomatology. Fourteen controls were tested with Modules 1 or 2 and four underwent the toddler module of the ADOS-2. All TD participants had a minimal severity score of 1, except one child who had a score of 3.

**Table 1.**
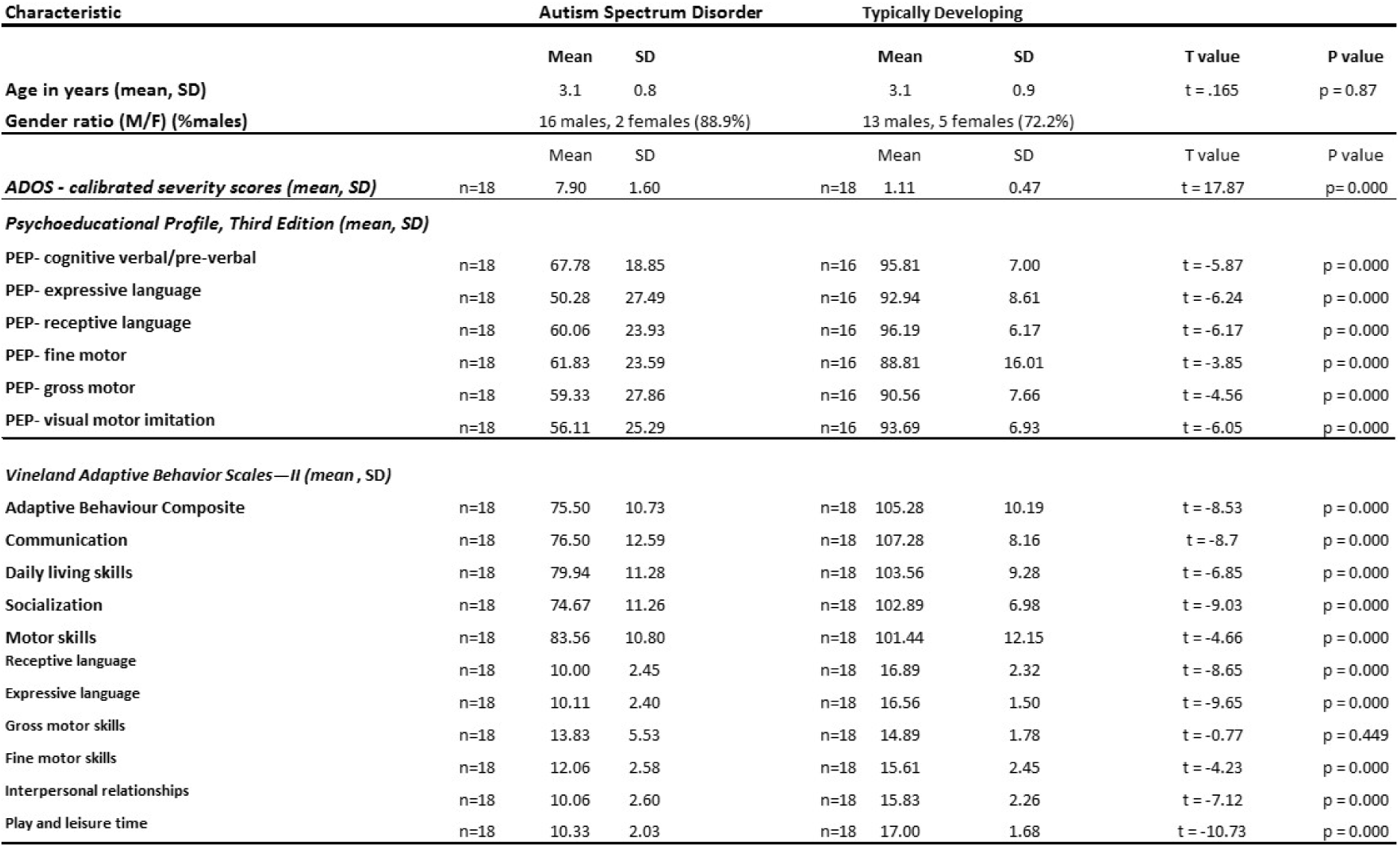
Characteristics of Study Participants

### Stimuli

Stimuli consisted of two video sequences of dynamic social images without audio information of approximatively two minutes each. These videos included ecologically valid and complex naturalistic dynamic images where young children practised yoga alone, imitated animal-like behaviours (behaving like a monkey or jumping like a frog), waived their arms, struck a pose, jumped, made faces or whistled (Yoga Kids 3; Gaiam, Boulder, Colorado, http://www.gaiam.com, created by Marsha Wenig, http://yogakids.com/). Presentation and timing of stimuli were controlled by Tobii Studio software (Sweden, http://www.tobii.com).

### Procedure and task

The experiment was conducted in a lit room at the office Médico-Pédagogique in Geneva. To familiarize the child with the procedure, the families received a kit containing a custom-made EEG replica cap and pictures illustrating the protocol in order to familiarize the children with the experiment two weeks prior to their first visit. Participants were seated on their parents lap in order to make them feel as secure as possible and to minimize head and body movements or alone. Once seated, the experimenter measured the circumference of the head and placed the corresponding cap on the participant’s head. A couple of minutes were taken in order to allow the participants to settle into the experiment’s environment and get used to the cap before starting the experiment. Following this, a five point eye-tracking calibration procedure was initiated using the Tobii system (Sweden, http://www.tobii.com). An attractive colourful object (either a kitten, a bus, a duck, a dog or a toy) was presented together with its corresponding sound on a white background and the participants had to follow the object visually. The recording and presentation of the visual stimuli started when a minimum of four calibration points were acquired for each eye. To best capture the child’s attention, we first showed them an age-appropriate animated cartoon, followed by some fractals and another animated cartoon. The block ended with a film containing dynamic social images, the condition of interest in the present experiment. All participants were presented with the same visual stimuli in the same order. Following the first block, impedances were rechecked and electrodes were readjusted where needed to maintain them below 40 k*Ohm*. A second block was then acquired (animated cartoon; animated fractals; animated cartoon; second condition of interest: dynamic social images). The experimenter continuously monitored the eye-tracking to ensure children were looking at the screen. The whole experiment lasted about half an hour. We used stringent criteria and only participants with the highest data quality were kept for subsequent analysis.

### Eye-tracking measurements

Eye-tracking data were recorded with the TX300 Tobii eye-tracking system (sampling rate resolution of 300 Hz). In order to analyse and quantify differences in visual exploration in our sample, we developed a data-driven method to define dynamic norms of the exploration of the visual scenes (Kojovic et al., in preparation). First, we applied a kernel density distribution estimation (Botev et al., 2010) on the eye-tracking data recorded from the TD group at each time frame of the films containing dynamic complex social images to compute a normative gaze distribution pattern. Then, for each of the participants with ASD individually, we computed a deviation index from this normative gaze distribution, and this, for each single time frame separately (Figure 5). We averaged these values across the two films to obtain a mean Proximity Index (PI) value. This index describes for a given ASD participant, his distance from the normative gaze distribution pattern calculated on the TD group. A high index value indicates a visual behaviour approaching the visual exploration of the TD participants (more similarity), while a low index indicates a visual behaviour deviating from the TD group (more dissimilarity).

**Figure 5.**
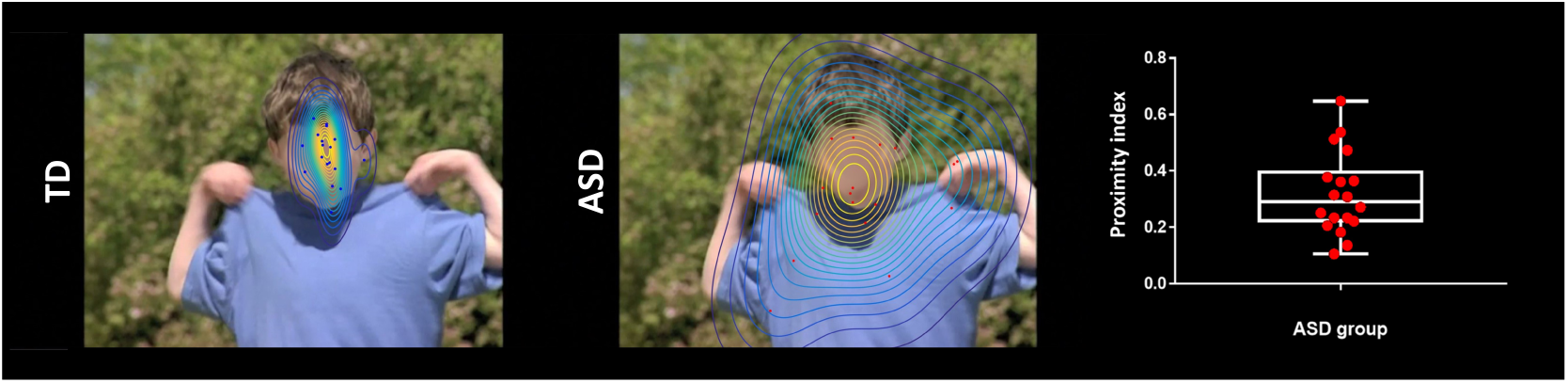
Exemplar single time frame of the normative gaze pattern for each group on one random time frame. Each dot represents the gaze position for an individual participant.

### EEG acquisition and preprocessing

The EEG was acquired with a Hydrocel Geodesic Sensor Net (HCGSN, Electrical Geodesics, USA) with 129 scalp electrodes at a sampling frequency of 1000*Hz*. On-line recording was band-pass filtered at 0–100*Hz* using the vertex as reference. Data pre-processing was done using Matlab (Natick, MA) and Cartool (http://sites.google.com/site/cartoolcommunity/). We down-sampled the montage to a 111-channel electrode array to exclude electrodes on the cheek and the neck since those are often contaminated with artefacts. Data were filtered between 1 and 40*Hz* (using non-causal filtering) and a 50*Hz* notch filter was applied. Each file was then visually inspected by one of the three EEG experts (HFS, TR, and RKJ) to exclude periods of movements artefacts. Periods where subjects were not looking at the screen were excluded. Independent component analysis (ICA) was performed on the data to identify and remove the components related to eye movement artefacts (eye blinks, saccades). Subsequently, channels with substantial noise were interpolated using spherical spline interpolation for each recording. Finally, the cleaned data were down-sampled to 125*Hz*, recalculated against the average reference and inspected by two EEG experts (HFS and AC) to ensure that no artefacts had been missed. One hundred and twenty artefact-free epochs of 1 second per participant were included for further analysis and were considered as a minimum to ensure high enough data quality.

### Electrical Source Imaging and selection of Regions of Interest

The general analysis strategy is summarized in Figure 6. Electrical source imaging (ESI) was performed to reconstruct the sources of brain activity that gave rise to the scalp EEG field. For this, we used an infant template head model (33-44 month) (using the Montreal Neurological Institute (MNI) brain) with consideration of skull thickness (Locally Spherical Model with Anatomical Constraints, LSMAC). 4159 solution points were equally distributed in the grey matter. We used a distributed linear inverse solution (Low Resolution Electromagnetic Tomography, LORETA (Pascual-Marqui et al., 1994)) to compute the 3-dimensional (3D) current source densities. We then projected this 3D dipole time-series, onto the predominant dipole direction of each region of interest (ROI) across time and epochs, therefore obtaining a scalar time-series series (Coito et al., 2016a, 2015; Plomp et al., 2015; Coito et al., 2016b). We parcelled the grey matter in 82 ROIs based on the automated anatomical labelling (AAL) digital atlas (Tzourio-Mazoyer et al., 2002), after normalization to the MNI space using SPM8 (Wellcome Trust Centre for Neuroimaging, University College London, UK, www.fil.ion.ucl.ac.uk/spm). In order to reduce the dimensionality of the solution space, we considered the solution point closest to the centroid of each ROI as representative of the source activity in that ROI for further analysis. This allowed us to obtain the source activity across time of 82 solution points, representative of 82 ROIs (Coito et al., 2016b).

**Figure 6.**
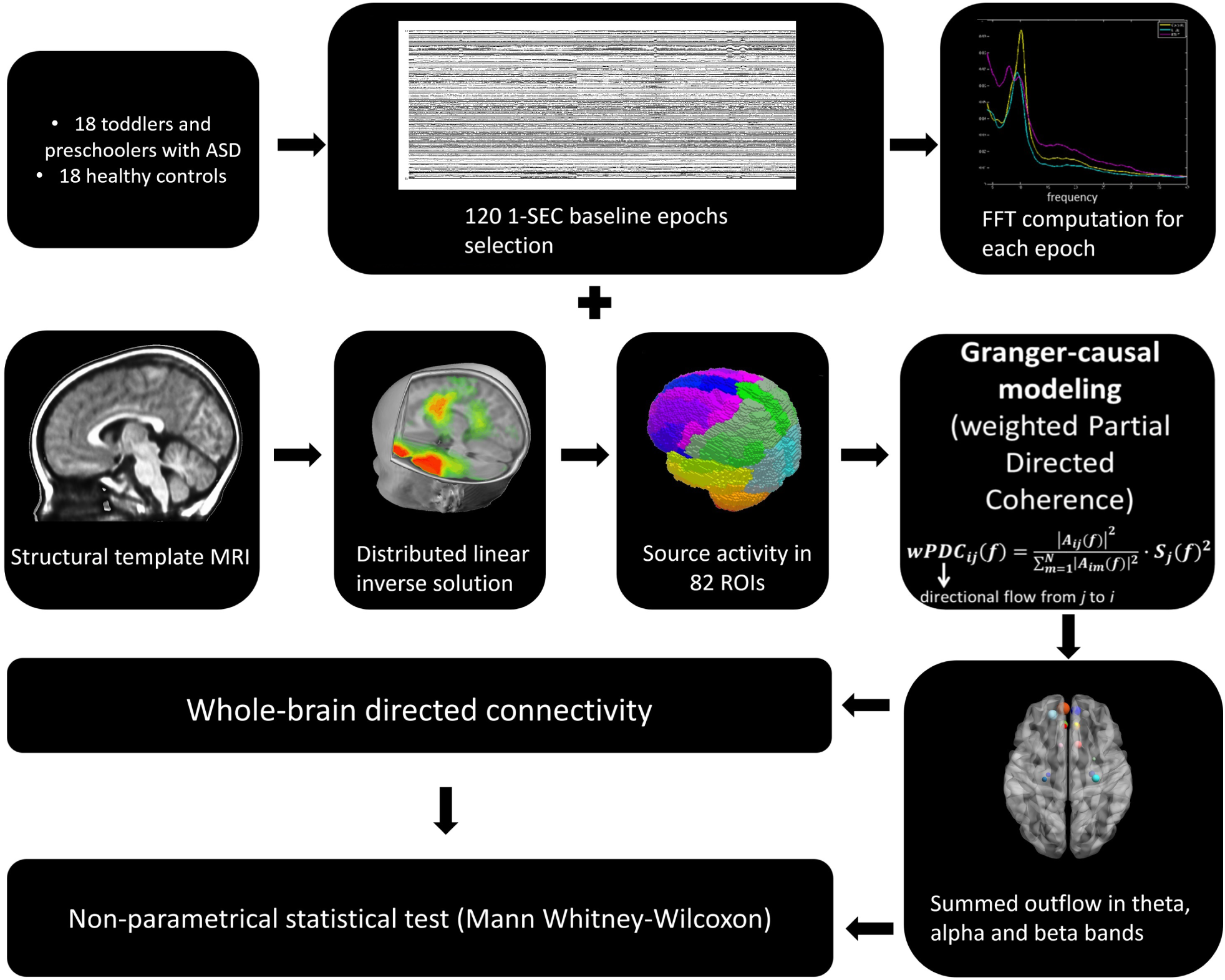
The general analysis strategy.

**Figure 7.**
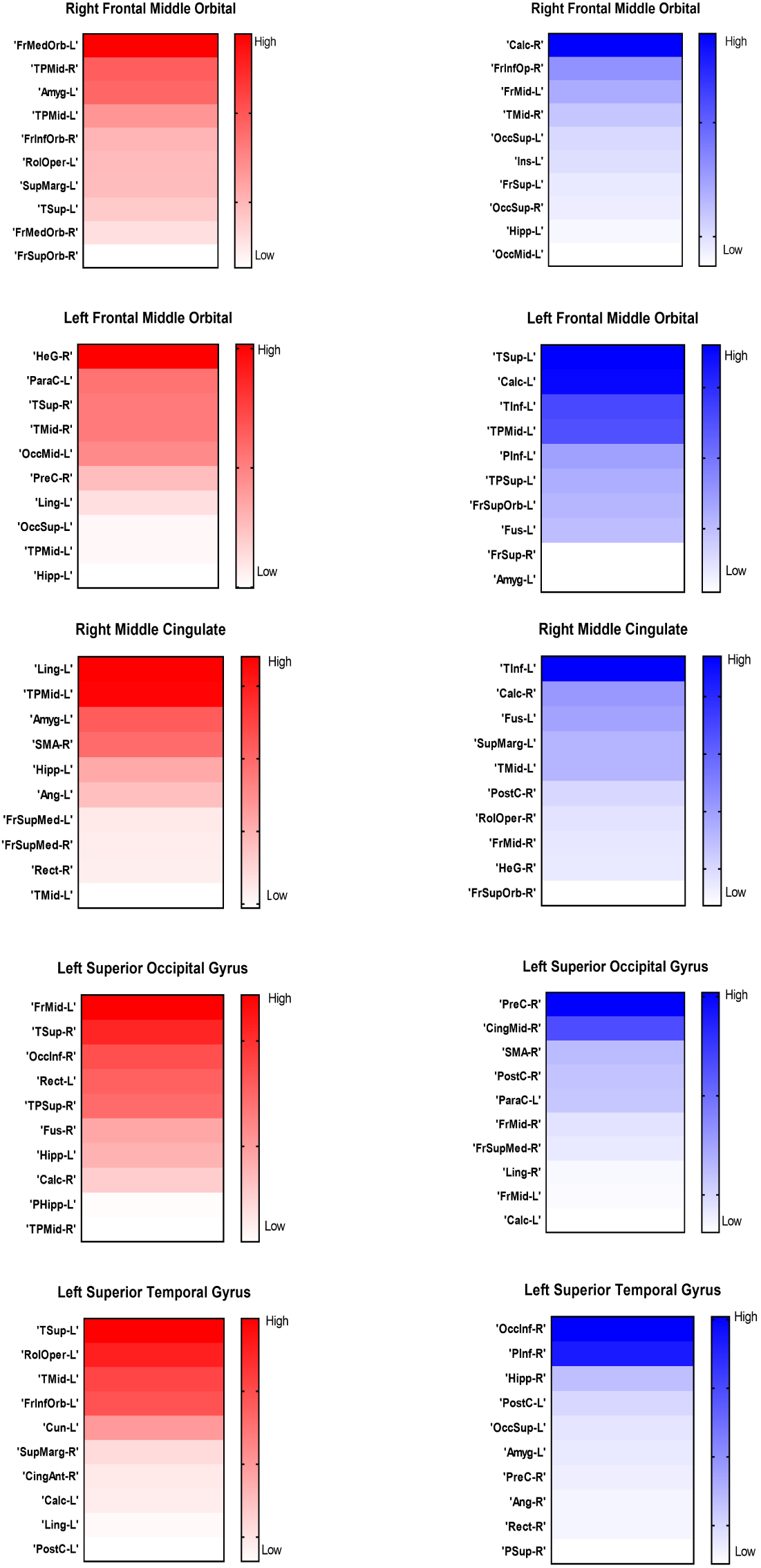
The directed connections from the five remaining ROIs : the left orbital part of the middle frontal gyrus, the right middle cingulate gyrus, the left superior occipital gyrus and the left superior temporal gyrus

### Directed Functional Connectivity using Granger-causality

Directed functional connectivity estimates the causal influence that one signal exerts onto another, facilitating the study of directional relationships between brain regions. It is commonly assessed using the concept of Granger-causality: given two signals in a process, if the knowledge of the past of one allows a better prediction of the presence of the other signal in the process, then the former signal is said to Granger-cause the latter signal (Granger, 1969). In order to estimate these directional relationships, we computed the weighted Partial Directed Coherence (*wPDC*) (Baccala & Sameshima, 2001; Astolfi et al., 2006; Plomp et al., 2014) using the 82 source signals. *PDC* is a multivariate approach, which considers all signals simultaneously in the same model and estimates brain connectivity in the frequency domain. It is computed using multivariate autoregressive models of a certain model order. Here, we used a model order of 5, corresponding to 40*ms*. The *wPDC* was computed for each subject and epoch and then the average of the *PDC* values within subjects was taken (Coito et al., 2016b). The average *PDC* was then scaled (0 - 1) across ROIs and frequencies (1 - 40*Hz*) by subtracting the minimum power and dividing by the range. In order to weight the *PDC* by the spectral power (*SP*) of each source signal, while avoiding frequency doubling, we computed the Fast Fourier Transform (*FFT*) for each electrode, applied ESI to the real and imaginary part of the *FFT* separately and then combined them (Coito et al., 2016a, 2015; Plomp et al., 2015; Yuan et al., 2008). The mean SP was obtained for each subject and scaled (that is 0-1, in the same way as *PDC*) for further details on the methodological approach to compute directed functional connectivity from electrical-source imaging signals, we refer the reader to (Coito et al., 2016b). For each subject, we obtained a 3D connectivity matrix (ROIs x ROIs x frequency), representing the outflow from one ROI to another for each frequency. For further analysis, we reduced the connectivity matrix to 3 frequency bands: theta (4 - 7*Hz*), alpha (7 - 12*Hz*) and beta (12 - 30*Hz*), by calculating the mean connectivity value in each band. For each subject and frequency band, we computed the summed outflow, which is the sum of the outflows (*wPDC* values) from a given ROI to all the others and reflects the driving importance of this ROI in the network: ROIs with high summed outflow strongly drive the activity of other ROIs. We identified the highest information transfer (summed outflow) in the theta band. Therefore, we focused our subsequent analysis on this frequency band. We carried out statistical comparisons of the summed outflows between subject groups using a non-parametrical statistical test (*Mann - W hitney - Wilcoxon,two - tailed, p <* 0.05). We then investigated the outflows from the ROIs that showed statistically significant summed outflow between groups to the whole brain (remaining 81 ROIs) and carried out a statistical comparison of these outflows between groups(*Mann - W hitney - Wilcoxon,two - tailed, p <* 0.05*,Ben jamini - Hochberg* = 0.05). We correlated (*Spearman - rho,two - tailed, p <* 0.05) the summed outflow results obtained in each of the 82 ROIs with ADOS-CSS scores, with developmental scores obtained from the PEP-3, with adaptive scores obtained from VABS-II and with the PI values obtained from the eye-tracking data. In all cases, correlation p-values were BenjaminiHochberg–corrected for multiple testing with *p* = 0.05. Connectivity computations were performed in Matlab. Figures 1,2,3 and 4 were produced using the BrainNet Viewer toolbox (Xia et al., 2013).

## Data availability

The data and codes that support the findings of the present work are available upon reasonable request to the corresponding authors Holger Franz Sperdin or Marie Schaer.

## Acknowledgements

The authors would like to express their gratitude to all the families who took part in this research. This research is supported by a grant from the National Centre of Competence in Research (NCCR) “SYNAPSY-The Synaptic Bases of Mental Diseases “ financed by the Swiss National Science Foundation (number: 51AU40-125759) and by private funding by the Fondation Pole Autisme (http://www.pole-autisme.ch). This work was further supported by a SNSF grant to M.S. (number: 163859), as well as SNSF grants number 320030-159705 to C.M.M and number 169198 and CRSII5170873 to S.V..R. K. J received individual support from a Marie Curie fellowship, which received funding from the European Union Seventh Framework Programme (FP7:2007-2013) under grant agreement number 267171.

## Author contributions

Conception and design of the experiment: H.F.S, A.C., S.E., C.M.M, G.P., T.R and M.S..Acquisition of data H.F.S., T.R,N.K, R.J, and M.F.. Analysis and/or interpretation of data: H.F.S, A.C. and M.S.. Drafting of the manuscript: H.F.S. and A. C.. All authors revised the manuscript critically for important intellectual content.

